# Uncovering the links between physical activity and prosocial behaviour: A functional near-infrared spectroscopy hyperscanning study on brain connectivity and synchrony

**DOI:** 10.1101/2023.08.18.553167

**Authors:** Toru Ishihara, Shinnosuke Hashimoto, Natsuki Tamba, Kazuki Hyodo, Tetsuya Matsuda, Haruto Takagishi

## Abstract

The prevalence of sedentary lifestyles in modern society raises concerns about their potential association with poor brain health, particularly in the lateral prefrontal cortex (LPFC), which is crucial for human prosocial behaviour. Here, we show the relationship between physical activity and prosocial behaviour, focusing on potential neural markers, including intra-brain functional connectivity and inter-brain synchrony in the LPFC. Forty participants, each paired with a stranger, underwent evaluation of neural activity in the LPFC using functional near-infrared spectroscopy hyperscanning during eye-to-eye contact and an economic game. Results showed that individuals with exercise habits and more leisure-time physical activity demonstrated greater reciprocity, less trust, longer decision-making time, and stronger intra-brain connectivity in the dorsal LPFC and inter-brain synchrony in the ventral LPFC. Our findings suggest that a sedentary lifestyle may alter human prosocial behaviour by impairing adaptable prosocial decision-making in response to social factors through altered intra-brain functional connectivity and inter-brain synchrony.

---

Prosocial behaviour encompasses actions that are generally considered beneficial to other people,^1^ such as volunteerism, charitable donations, fair resource distribution, cooperative behaviour, and reciprocity of favours. It plays a critical role in an individual’s life. Its significance is evident in various scenarios, including natural disasters, such as earthquakes and tsunamis, during which mutual aid becomes a crucial survival strategy. Beyond these practical examples, human prosocial behaviour has been linked to labour market success, irrespective of the continent or country’s economic development, as individuals exhibit higher income and employment rates worldwide.^2^ Moreover, engaging in prosocial behaviour has been causally associated with improved health outcomes,^3,4^ including enhanced psychological well-being^5^ and physical health,^6^ while serving as a protective factor against the detrimental effects of stress^7^ and reducing stress-related mortality risk.^8^ Notably, human prosocial behaviour is influenced by a combination of genetic and environmental factors, with heritability accounting for only approximately 20%–30% of the observed variance.^9,10^ Consequently, identifying the environmental factors that can modulate human prosocial behaviour is crucial from both economic and health perspectives.

A sedentary lifestyle is a factor that can hinder human prosocial behaviour. The prevalence of sedentary lifestyles has become a significant global health concern, with approximately 81% of adolescents and 27.5% of adults worldwide failing to meet recommended levels of physical activity.^11,12^ A sedentary lifestyle is associated with an elevated risk of various diseases and mortality.^13,14^ Moreover, research conducted over the past three decades has underscored the crucial role of physical activity in maintaining and promoting brain and cognitive health.^15–17^ Among the brain regions affected by physical activity, the lateral prefrontal cortex (LPFC) and its function have received considerable attention in the literature.^15–17^ Given that the LPFC is considered one of the primary neural bases of human prosocial behaviour,^18–20^ there is a potential concern that global trends towards a sedentary lifestyle may have an adverse association with human prosocial behaviour.

Existing evidence on the relationship between physical activity and prosocial behaviour suggest both positive^21–23^ and null or negligible associations.^24–26^ Consequently, there is a lack of clear evidence showing the effect of physical activity on human prosocial behaviour. A possible explanation for the discrepancies observed across these studies is the use of questionnaire-based, subjectively measured prosocial behaviour, which introduces various biases, such as social desirability, recall, and reference biases. Further, prosocial behaviour measured subjectively by questionnaires only provides a rough reflection of an individual’s prosocial tendencies and does not assess specific aspects such as trust and reciprocity. Assuming that the LPFC serves as a neural foundation connecting physical activity and prosocial behaviour, it is difficult to accurately capture the impact of physical activity without evaluating the LPFC-dependent domains of prosocial behaviour. By observing actual behaviour in near-real-world economic exchanges and investigating the functions of the LPFC during prosocial decision-making processes, we can enhance existing knowledge and gain valuable insights into the neural mechanisms underlying the link between physical activity and prosocial behaviour.

This study aimed to investigate the relationship between physical activity and objectively measured LPFC-dependent prosocial behaviour, and the contribution of LPFC functioning during prosocial decision-making processes. We employed the trust game,^27^ a well-known economic game, as a measure to evaluate prosocial behaviour. In this game, participants are given tasks that involve transferring a certain amount of money to a trustee; the transferred amount is typically multiplied by a factor (often three). The trustee then decides the amount of money to return, ultimately determining the payoff for both players. The trust game assesses two critical dimensions of prosocial behaviour: trust and reciprocity. Trust refers to the extent to which individuals are willing to trust strangers, particularly in situations where they must rely on the actions or decisions of others. Reciprocity pertains to the degree to which individuals reciprocate or respond to cooperative behaviour when they are trusted by others. The trust game was selected as an appropriate measure to evaluate prosocial behaviour because it is associated with LPFC-dependent, strategically motivated prosocial acts.^20^ Unlike self-report questionnaires, the trust game allows for the assessment of actual LPFC-dependent prosocial decision-making in the context of social interaction.

This study focused on two systems within the LPFC that could serve as neural markers for the interaction between physical activity and human prosocial behaviour. The first system is cognitive control, which involves coordination of mental processes and actions according to current goals and future plans; this is primarily regulated by the dorsal LPFC (DLPFC).^28^ Humans demonstrate adaptable prosocial decision-making in response to social norms, including fairness norms and other social factors, by controlling for and overriding intuitive and/or preferential choices.^29–33^ Cognitive control is believed to be crucial for facilitation of flexible behavioural adaptations.^18–20^ Cognitive control and the DLPFC play prominent roles in complying with social norms, including those related to fairness.^34,35^ The causal association of DLPFC function with prosocial behaviour has been demonstrated as decreased and increased prosocial decisions through cathodal (inhibitory) and anodal (excitatory) transcranial direct current stimulation (tDCS), respectively, over the DLPFC.^34^ Considering that physical activity has been shown to be associated with greater cognitive control and DLPFC function,^15–17^ it is plausible to hypothesise that physical activity is positively associated with prosocial behaviour.

The second LPFC system of interest in this study was inter-brain synchrony, which involves neural oscillations within specific frequency bands that synchronise across the brains of individuals engaged in social interactions.^36^ The proposed neural mechanism is believed to enhance social interaction by facilitating the integration of multiple brains.^36^ A meta-analysis of previous studies provided evidence of statistically significant inter-brain synchrony during cooperation,^37^ with significant synchrony in the LPFC.^37^ Inter-brain synchrony has also been observed during simple eye-to-eye contact^38,39^ and economic exchange,^40,41^ and the strength of synchrony has been linked to increased cooperative choices.^40^ While the exact factors that elicit inter-brain synchrony are still under discussion,^42^ this phenomenon has been robustly observed and proposed as one of the neural bases of human prosocial behaviour.^41^ We assessed inter-brain synchrony during eye-to-eye contact and a trust game as possible neural markers of the interaction between physical activity and prosocial behaviour. Although direct evidence of the potential link between physical activity and inter-brain synchrony is currently lacking, it is possible that physical activity influences the ability to synchronise with the brains of others, given its association with LPFC structure and function.^43,44^

In this study, we investigated the functions of the LPFC and prosocial behaviour in relation to physical activity by comparing naturalistic face-to-face interactions to those with a fictitious person. This comparison allowed us to assess the contributions of inter-brain synchrony. Previous research has demonstrated that engaging in tasks simultaneously with a partner, such as looking at a face and engaging in prosocial behaviour, can lead to similar neural responses and induce inter-brain synchrony.^42^ By comparing these two conditions in which participants engage in the same tasks, we can distinguish the synchronization that occurs when individuals work on the same tasks from the brain synchrony induced by face-to-face interaction, and determine its contribution of inter-brain synchrony to the interaction between physical activity and prosocial behaviour. Furthermore, face-to-face communication has been demonstrated to make social norms more salient^45^ and increase prosocial behaviour compared to non-face-to-face interactions, such as those with human-like avatars.^46,47^ This suggests that cognitive control plays a role in facilitating behavioural adaptations during face-to-face interactions. Therefore, we hypothesised that the associations between physical activity, cognitive control, inter-brain synchrony, and prosocial behaviour would be more pronounced in the context of face-to-face communication.

Functional near-infrared spectroscopy (fNIRS) was used to assess LPFC function during eye-to-eye contact and the trust game. This technique is advantageous over conventional methods such as electroencephalography or functional magnetic resonance imaging (fMRI) during face-to-face naturalistic communication because of its reduced susceptibility to movement artefacts and body restrictions. To assess LPFC function and inter-brain synchrony simultaneously, we evaluated intra- and inter-brain functional connectivity during eye-to-eye contact and the trust game. Functional connectivity refers to the temporal correlation of activation that occurs at rest or during engagement in tasks among different brain regions.^48^ Compared with event-related designs, this approach allowed us to free participants from time restrictions, making it more suitable for the naturalistic nature of the present study design. Studies have shown that intra-brain functional connectivity in the LPFC is elevated with increased brain workload and task difficulty requiring cognitive control,^49^ and is positively associated with task performance.^50^ Therefore, engagement in cognitive control and its function during social interactions is reflected in the observed intra-brain functional connectivity in the LPFC. In addition, we assessed the resting-state intra-brain functional connectivity which can be a neural marker of the link between physical activity and prosocial behaviour. Resting-state functional connectivity reflects cognitive control capacity and is positively associated with both cognitive control^51^ and prosocial behaviour.^52^ Some evidence suggests that inter-hemispheric connectivity is strongly associated with physical activity, cognitive control, and prosocial behaviour.^51–54^ We evaluated inter- and intra-hemispheric connectivity separately. To assess inter-brain synchrony, which refers to the functional connectivity between two brains, we utilised the hyperscanning technique; this is an experimental approach that enables the simultaneous recording of brain activities of two individuals. We hypothesised that there is a positive association between physical activity and trust, reciprocity, intra-brain functional connectivity, and inter-brain synchrony, and that this association can be pronounced face-to-face relative to face stimulus conditions.

## Results

A schematic overview of the study is presented in Fig. 1. Here, 40 participants completed two conditions in a randomised order after resting-state functional connectivity evaluation: 1) face-to-face condition—eye-to-eye contact with their partner, and 2) face stimulus condition—eye-to-eye contact with the face stimulus of a fictitious person displayed on the screen (Fig. 1a). Following each condition, participants played trust games with their partner or the fictitious person displayed on the screen (Fig. 1a). Neural activity in the LPFC was recorded using fNIRS hyperscanning (Fig. 1a and b). The fNIRS device consisted of 16 active channels and the probe sets were symmetrically positioned on the scalp over the prefrontal regions of the participants (Fig. 1c). To evaluate intra-brain functional connectivity and inter-brain synchrony during rest, eye-to-eye contact, and the trust game, two sets of time-series wavelet coherences were computed (Fig. 1d and e). After the experiments, 38 participants reported that they had not known each other before the experiment, while one participant reported that he had seen his partner before. Another participant reported that he and his partner were acquaintances, whereas the partner reported that they knew nothing of each other. Additional interviews revealed that they had been in the same group during job interviews and did not have any other interactions previously.

**Fig. 1.**
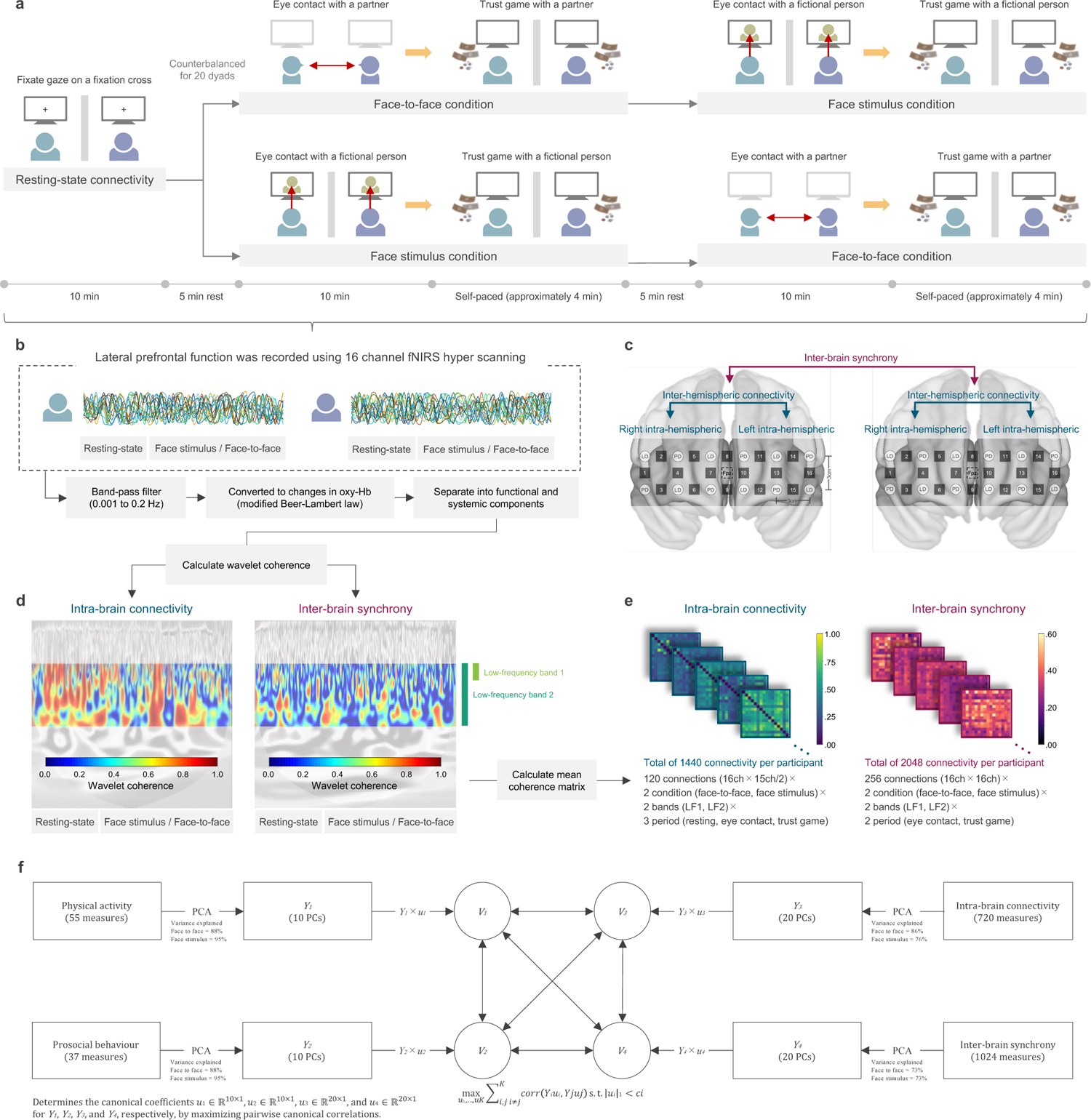
Schematic Overview of the Study. a. A schematic diagram illustrating the experimental conditions and depicting the experimental design. b. Pipeline for fNIRS analysis. c. Schematic diagrams showing the placement of fNIRS channel montage and the inter- and intra-brain connectivity. LD = laser diode emitter; PD = photodiode detector. d and e. Calculation procedure for intra-brain connectivity and inter-brain synchrony, along with representative wavelet coherence maps (left panel: inter-brain synchrony, right panel: intra-brain connectivity). LF1 = low-frequency band 1 (LF1: 0.015 to 0.15 Hz), LF2 = low-frequency band 2 (LF2: 0.08 to 0.15 Hz). f. Analysis schematic depicting the sparse multiset canonical correlation analysis.

### Face-to-face versus face stimulus condition on prosocial behaviour

The results of the two-way ANOVA on reciprocity showed significant main effects of the condition (*F* = 9.28, *p* = .004, partial η^2^ = .19; Fig. 2a and b). Additionally, there were significant main effects of the amount of money received (*F* = 47.54, *p* < .001, partial η^2^ = .55; Fig. 2a). However, no credible main effects of the initial choice, final decision, or interactions were detected (Table S1). There were no credible differences in the trust and decision time measures between the conditions (Table S1).

**Fig. 2.**
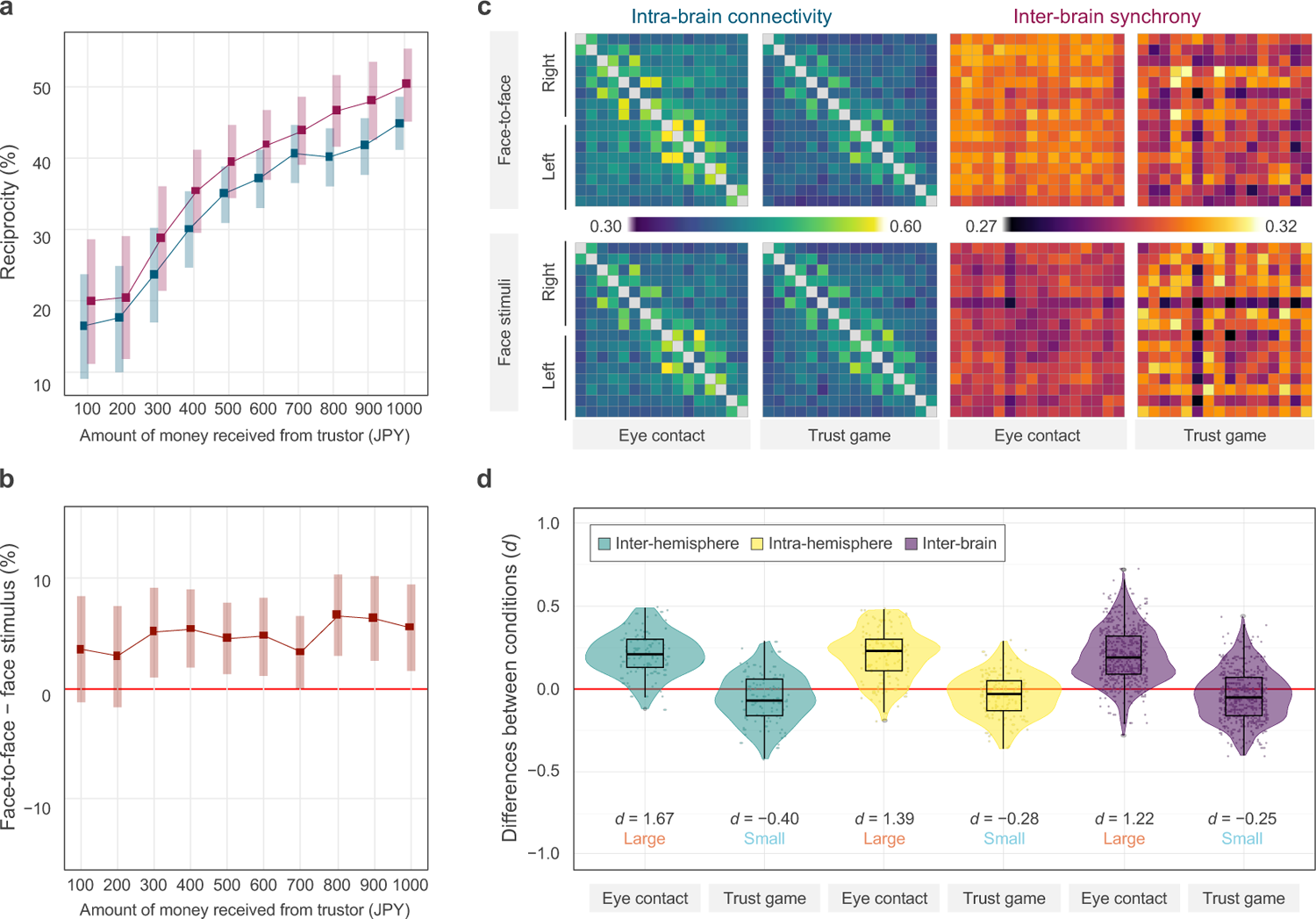
Prosocial behaviour, intra-brain functional connectivity, and inter-brain synchrony and their differences between face-to-face and face stimulus conditions. a. Mean reciprocity in each condition. Shaded bars represent 95% confidence intervals. b. Mean differences in reciprocity between conditions. Shaded bars represent 95% confidence intervals. c. Mean coherence matrix in each combination of channels. The channels are arranged in the order from channel 1 to channel 16, moving from left (top) to right (bottom) of the matrix. For the intra-brain functional connectivity, the upper left and the lower right quarters of the matrix represent intra-hemispheric functional connectivity, while the upper right and the lower left quarters represent inter-hemispheric functional connectivity. Functional connectivity including channels 8 and 9 are considered part of the intra-hemispheric connectivity. d. Comparisons of intra-brain functional connectivity and inter-brain synchrony across conditions. Each plot represents the effect size (Cohen’s *d*) for the differences in intra-brain functional connectivity and inter-brain synchrony between conditions across each combination of channels. Cohen’s *d* presented in the graph represents the size of the effects for the difference from zero.

### Face-to-face versus face stimulus condition on intra-brain functional connectivity and inter-brain synchrony

During eye contact, the face-to-face condition exhibited greater inter- and intra-hemispheric functional connectivity (Cohen’s *d*s for the difference from zero = 1.67 and 1.39, respectively; Fig. 2c and d) and inter-brain synchrony (Cohen’s *d* for the difference from zero = 1.22; Fig. 2c and d) compared with the face stimulus condition, while no such credible differences were found during the trust game (Cohen’s *d*s for the difference from zero = −0.40 to −0.25; Fig. 2c and d).

### Relation of physical activity to prosocial behaviour and brain function

Physical activity data were collected using questionnaires,^54–56^ which gathered information on various types of physical activities, including regular exercise, job-related activities, transportation, domestic chores, gardening, leisure activities, and sedentary behaviour. In this study, we obtained a comprehensive dataset comprising the following number of measures: physical activity, 55; prosocial behaviour, 37; intra-brain connectivity, 720; and inter-brain synchrony, 1024. A list of physical activity and prosocial behaviour measures is provided in Table S2.

To investigate the relationships among these four sets of variables—physical activity, prosocial behaviour, intra-brain functional connectivity, and inter-brain synchrony—we employed the sparse multiset canonical correlation analysis (SMCCA; Fig. 1f), which is specifically designed to identify multivariate associations among more than three modalities, extending the conventional canonical correlation analysis.^57^ By applying SMCCA, we estimated canonical variates that capture shared covariance patterns across participants in the four variables. To ensure a well-posed solution and avoid overfitting and a rank-deficient solution, we performed dimensionality reduction using principal component analysis (PCA) prior to application of SMCCA (Fig. 1f). We employed a non-parametric permutation test to assess the statistical significance of the SMCCA components.

In the face-to-face condition, the results revealed a single significant SMCCA mode that related physical activity to prosocial behaviour, inter-brain synchrony, and intra-brain functional connectivity (sum of correlation coefficients = 5.14, *p* = 0.008; Fig. 3a and b), whereas no credible evidence of a significant mode was revealed in the face stimulus condition (sum of correlation coefficients = 2.12, *p* = 0.28; Fig. S1a). When inter-brain synchrony was excluded from the SMCCA of the face stimulus condition, no credible evidence of a significant mode was found (sum of correlation coefficients = 1.05, *p* = 0.50; Fig. S1b). We examined the physical activity and prosocial behaviour measures strongly associated with the identified canonical variates (Fig. 3c). Regular exercise and leisure-time physical activity had large positive loadings (Fig. 3c). Furthermore, reciprocity measures had larger positive loadings; the greater the amount of money received from the trustor, the higher was the trend (Fig. 3c). However, trust measures had negative loadings (Fig. 3c). The decision time measures tended to positively correlate with the loadings, except for reciprocity, when receiving 200 and 300 JPY (Fig. 3c). The canonical loadings of prosocial behaviour were positively associated with the effect sizes of their difference between conditions (Spearman’s ρ = 0.32 to 0.68, Fig. S2).

**Fig. 3.**
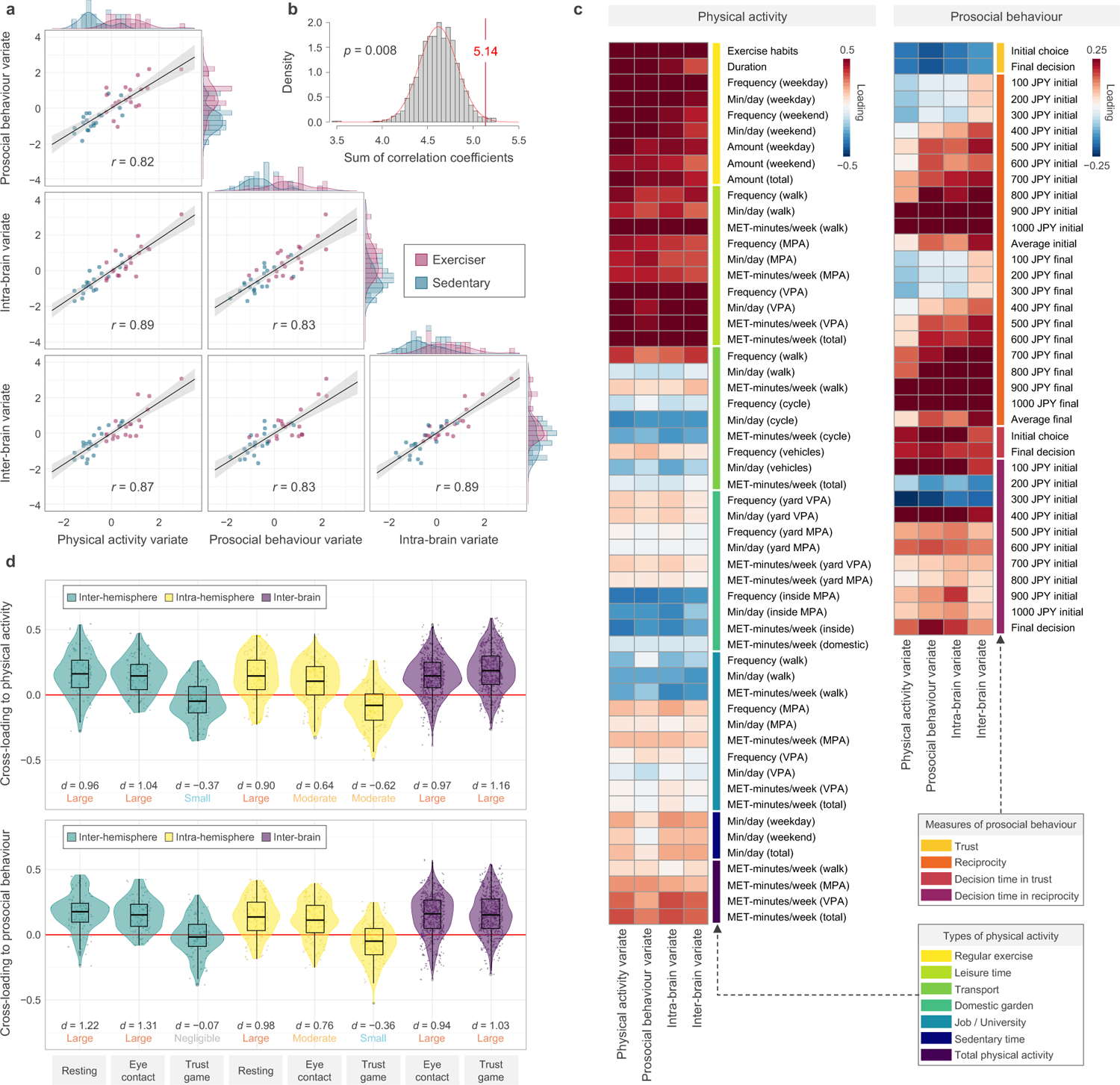
Results of SMCCA during the face-to-face condition. a. The principal SMCCA mode, represented as a scatter plot of physical activity, prosocial behaviour, intra-brain functional connectivity, and inter-brain synchrony variates, with one point per participant. An example of a physical activity measure (exercise habits) is indicated by different colours. b. Results of the non-parametric permutation test with 1000 iterations. c. The correlation of physical activity and prosocial behaviour measures with the identified variates. d. The correlation of intra-brain functional connectivity and inter-brain synchrony measures with physical activity (upper panel) and prosocial behaviour (lower panel) variates. Cohen’s *d* presented in the graph represents the size of the effects for the difference from zero.

Next, we examined the intra-brain connectivity and inter-brain synchrony measures that were strongly associated with the identified physical activity and prosocial behaviour variates (Fig. 3d). The average cross-loadings of intra-brain connectivity measures at rest and during eye contact with physical activity and prosocial behaviour variates were significantly different from zero with moderate to large effect sizes (Cohen’s *d*s = 0.64 to 1.31; Fig. 3d), while no credible association was found during the trust game with negligible to moderate effect sizes (Cohen’s *d*s = −0.62 to −0.07; Fig. 3d). The average cross-loadings of inter-brain synchrony measures during eye contact and trust games with physical activity and prosocial behaviour variates were significantly different from zero, with large effect sizes (Cohen’s *d*s = 0.94 to 1.16; Fig. 3d).

To aid regional interpretations, the cross-loadings of intra-brain functional connectivity and inter-brain synchrony measures with physical activity and prosocial behaviour variates in each combination of channels (Fig. 4a) were averaged in each channel, and spatial maps were created (Fig. 4b). In terms of intra-brain functional connectivity, the DLPFC exhibited strong correlations with physical activity and prosocial behaviour variates (Fig. 4b), whereas for inter-brain synchrony, the ventrolateral prefrontal cortex (VLPFC) displayed strong correlations with physical activity and prosocial behaviour variates (Fig. 4b). Furthermore, the regional correlations of the average weights between intra-brain functional connectivity and inter-brain synchrony showed an inverse association (Fig. S3).

**Fig. 4.**
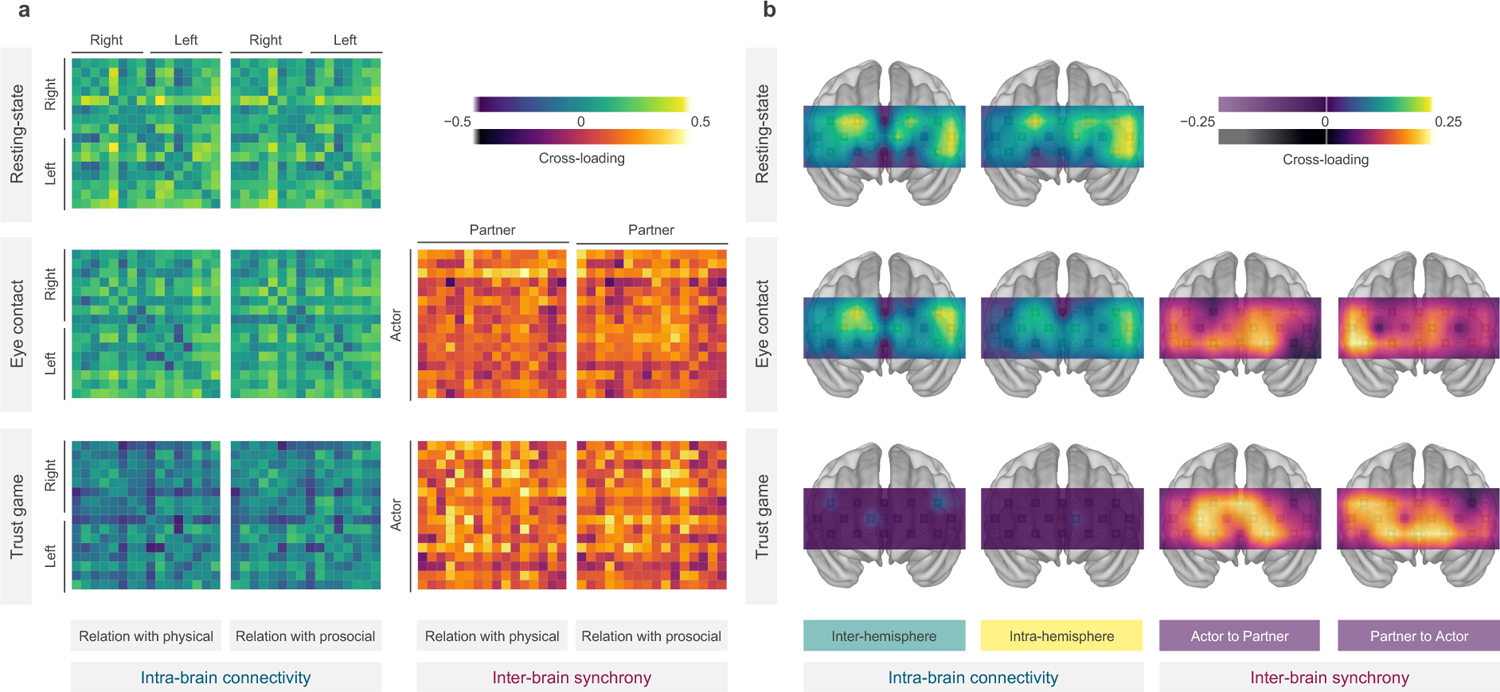
Canonical cross-loadings of intra-brain functional connectivity and inter-brain synchrony with physical activity and prosocial behaviour variates. a. Matrix displaying the cross-loadings for each combination of channels, which are arranged in sequential order from 1 to 16, progressing from left (top) to right (bottom) within the matrix. For the intra-brain functional connectivity, the upper left and lower right quarters of the matrix represent intra-hemispheric functional connectivity, while the upper right and lower left quarters represent inter-hemispheric functional connectivity. For inter-brain synchrony, the vertical direction lists the channels of the actor, while the horizontal direction lists the channels of the partner. b. SMCCA mode intra-brain functional connectivity and inter-brain synchrony average cross-loadings for physical activity and prosocial variates and associated spatial maps. The loadings are averaged because the cross-loading for physical activity and prosocial variates were highly correlated (Pearson’s r = 0.77 to 0.94).

### Sensitivity analyses

To confirm the robustness of SMCCA while varying the number of principal components, we repeated the entire statistical testing for a range of PCs for inter- and intra-brain connectivity data. The correlation among the canonical cross-loadings while varying the number of principal components showed almost no change in the results—the correlations were [0.63 to 0.98] (Fig. S4). To confirm the present results, which are affected by the participants’ social value orientation that reflects stable personality traits towards prosocial behaviour and sleepiness during fNIRS recording, a sensitivity analysis was performed. The inter-relationships between physical activity, prosocial behaviour, intra-brain functional connectivity, and inter-brain synchrony were fundamentally unchanged after controlling for these factors (sum of correlation coefficients = 5.07, *p* = 0.004; Fig. S5). We also tested the robustness of the present results by using deoxy-Hb data. The results of this analysis were identical to those of the original analysis (Fig. S6).

## Discussion

This study aimed to investigate the association between physical activity and prosocial behaviour as well as its neural markers during social interaction and prosocial decision-making. We found positive correlations between physical activity, reciprocity, decision-making time, intra-brain functional connectivity in the DLPFC, and inter-brain synchrony in the VLPFC in face-to-face situations. With trust, a negative correlation was found. These findings emphasise the potential association between the sedentary lifestyle pandemic and human prosocial behaviour, possibly due to alterations in the functioning of two prefrontal systems: intra-brain functional connectivity in the DLPFC and inter-brain synchrony in the VLPFC. Consequently, a sedentary lifestyle can be considered a contributing environmental factor that influences human prosocial behaviour.

The present findings offer insights into the mixed results of previous studies on the association between physical activity and prosocial behaviour.^21–26^ Our study revealed a positive correlation between physical activity and reciprocity as well as a negative correlation with trust. The results of reciprocity were in accordance with our a priori hypothesis, whereas those of trust were not. In a trust game, the trustors and trusted parties are differentially sensitive to risks and benefits.^58^ Trustors primarily focus on the risks associated with trusting, whereas trustees base their decisions on the level of benefits they have received.^58^ Specifically, a trusting tendency is more likely to prevail in cases of low risk; however, the likelihood of trusting does not depend on the level of benefit provided to the trusted parties.^58^ Conversely, reciprocity is more likely in case of greater benefit; however, it does not depend on the risk level faced by the trustor.^58^ Considering the distinct dynamics of reciprocity and trust, our study suggests that physical activity is associated with heightened altruism and cautious trust.

These interpretations are supported by the results of intra-brain functional connectivity. Physical activity was associated with stronger intra-brain functional connectivity in the DLPFC, which in turn was associated with increased reciprocity, decreased trust, and longer decision-making times. These findings suggest that cognitive control serves as a neural marker for the relationship between physical activity and reciprocity as well as cautious trust. Although previous studies have provided evidence that cognitive control and the DLPFC play a prominent role in complying with social norms^34,35^ and are associated with increased prosocial decisions,^34^ these associations have also been found to be heavily situation-dependent.^59^ Inhibitory tDCS over the DLPFC led to an increase in the rule-abiding capacity of the participants, even when the rules required losing money or causing financial harm to another person.^60^ However, excitatory tDCS led the participants to violate rules that conflicted with their freely chosen preferences.^60^ These findings suggest that DLPFC function does not simply increase or decrease prosocial behaviour; rather, it integrates the costs/risks and benefits of rules to align decisions with internal goals, ultimately enabling flexible adaptation of social behaviour.^60^ For example, previous studies have demonstrated that greater DLPFC thickness and longer decision-making times are associated with decreased prosocial behaviour^61^ and this trend is stronger in highly prosocial individuals.^30,31^ A possible explanation for this association is the risk of exploitation by others through cognitive control.^30^ In human society, engagement in prosocial behaviour can cause exploitation; therefore, intuitive engagement in prosocial behaviour in risky situations can be dangerous. Hence, it is necessary to carefully consider and calculate risks using cognitive control, and sometimes make selfish decisions. However, prosocial behaviour without inherent risks, such as reciprocity when the amount offered by the trustor is high, relies on social norm compliance, which requires cognitive control^34,35^ to suppress the desire to increase one’s own benefits. Individuals who engage in higher levels of physical activity might utilise cognitive control and behave non-cooperatively during engagement in trust behaviours involving risks, while behaving cooperatively during engagement in reciprocity involving fairness norms. A positive association between physical activity and prosocial behaviour was observed exclusively in the face-to-face condition, with no credible association found in the face stimulus condition.

This finding supports our a priori hypothesis that the positive relationship between physical activity and prosocial behaviour is more pronounced in real-life face-to-face situations than in non-face-to-face interactions or interactions with fictitious individuals. The amount of reciprocity exhibited in the face-to-face condition was significantly higher than that in the control condition, indicating that the participants might demonstrate adaptable prosocial decision-making in response to situational and social factors. These results support the previous knowledge that face-to-face communication makes social norms more salient^45^ and increases prosocial behaviour compared to non-face-to-face interactions.^46,47^ Furthermore, the effect sizes of the differences in behavioural measures between conditions during the trust game were positively correlated with the canonical loadings. These correlations suggest that behaviour during the trust game, which showed greater changes from facial stimuli to face-to-face conditions, is strongly associated with physical activity, intra-brain functional connectivity, and inter-brain synchrony. Thus, physical activity may be associated with reciprocity via fairness norm compliance using cognitive control, especially in face-to-face situations.

Additionally, the specific association between physical activity and prosocial behaviour in the face-to-face condition can be explained by inter-brain synchrony, which was positively associated with physical activity and reciprocity during eye-to-eye contact and the trust game. These findings are consistent with those of a previous study that demonstrated a link between the strength of synchrony and increased cooperative choices.^40^ In this study, no contribution from inter-brain synchronisation in the facial stimulation condition was observed. This could be due to the fact that the individuals measuring brain synchronization and those participating in the trust game were different. Therefore, it is reasonable to consider that the face-to-face condition would exhibit a stronger relationship between physical activity and reciprocity than would the facial stimulation condition as it considers the additional contribution of inter-brain synchronisation. Although numerous studies have demonstrated an association between physical activity and LPFC structure and function,^43,44^ none of them attempted to examine the association between physical activity and inter-brain synchrony. This study expands on previous knowledge by demonstrating the possible influence of physical activity on the ability to synchronise with the brains of others.

Interestingly, the regional correlations of the average weights between intra-brain functional connectivity and inter-brain synchrony showed an inverse association. These results indicate that two distinct LPFC systems contribute to the interaction between physical activity and prosocial behaviour. We found that inter-brain synchrony strongly contributed to the association between physical activity and prosocial behaviour in the VLPFC, which is a well-known region that plays a crucial role in voluntary emotion regulation, particularly in social interactions; this demonstrates its essential role in the downregulation of negative emotions and upregulation of positive emotions.^62^ Thus, inter-brain synchrony in this region could contribute to the sharing of an emotional state with a partner which, in turn, may increase reciprocity during the trust game.

However, the inverse association between inter-brain synchrony and trust observed in this study contradicts the results of a previous study which showed that the strength of synchrony is linked to increased cooperative choices.^40^ This discrepancy could be explained by differences in the cognitive processes between the sender and receiver roles during the trust game. Participants in the trust game exhibited clear social preferences when they assumed the role of the receiver.^63^ However, in the role of the sender, they failed to differentiate between a human partner and a non-social, lottery-like scenario in which a computer assumes the role of the receiver.^63^ Considering that the decision-making of the sender is somewhat independent of the partner and is significantly influenced by risk aversion, lack of a clear association with inter-brain synchronisation is reasonable.

The contribution of intra-brain functional connectivity was observed only during rest and eye-to-eye contact, and no credible contribution was detected during the trust game. The reason for the lack of correlation between intra-brain functional connectivity during the trust game and physical activity or prosocial behaviour remains unclear. It could be that the participants were initially informed to engage in the trust game with a partner or fictitious individual after eye-to-eye contact. Participants may have already made a decision during the eye-contact phase in the trust game. This could be explained by the absence of significant differences in intra-brain functional connectivity between the conditions during the trust game, despite significant differences observed during eye contact.

In terms of physical activity, regular exercise and leisure time, particularly those involving a high intensity, have shown a strong and positive association with prosocial behaviour, intra-brain functional connectivity, and inter-brain synchrony when compared to other types of physical activity. These findings could be explained by the relationship between the type of physical activity and cognitive control, as well as DLPFC function, along with social interaction during physical activity. Numerous studies have consistently reported a positive relationship between cardiorespiratory fitness and cognitive control, as well as DLPFC function,^15,16,64^ suggesting that physical activity at a sufficient intensity can effectively improve cardiorespiratory fitness and enhance cognitive control. Furthermore, regular exercise during leisure time is expected to involve more social interaction than does physical activity for transportation purpose or during domestic gardening, which may moderate the effects on prosocial behaviour. However, this hypothesis should be confirmed by future research that specifically focuses on the modality of physical activity and its impact on social interactions.

The present study has several limitations. First, due to the cross-sectional nature of the study, we could not establish a causal relationship between physical activity and prosocial behaviours. Personality traits related to prosocial behaviour may lead to increased physical activity, rather than vice versa. However, it is important to note that the main findings remain robust even after accounting for the confounding effects of social value orientation. This suggests that the observed association between physical activity and prosocial behaviour persists even after considering the stable personality traits related to prosocial behaviour and their impact on physical activity. Therefore, we believe that the likelihood of physical activity influencing human prosocial behaviour is high, particularly in terms of adaptable decision-making in response to situational and social factors. This argument is further supported by the lack of association between physical activity and prosocial behaviour in the face stimulus condition. To clarify the causal relationship between physical activity and prosocial behaviours, more rigorous study designs should be used in the future. Second, while we demonstrated a novel contribution of inter-brain synchrony to the covariation between physical activity and prosocial behaviour, the interpretation of inter-brain synchrony is still debated.^42^ The specific meaning of brain synchrony in this context and its contribution to prosocial behaviour should be examined in future studies. Third, our study focused primarily on the function of the LPFC; however, other regions involved in the empathy and mentalizing network, such as the temporal–parietal junction and posterior cingulate cortex, also contribute to human prosocial behaviour.^18^ Recent findings suggest that exercise can enhance social brain network function,^65,66^ thereby suggesting the role of these additional regions in this association. Future studies should incorporate other imaging methods, such as fMRI, to explore the involvement of these regions. Finally, the generalisability of our findings remains unclear. In future research, it will be important to test the replicability of the present results in other cultures and populations, such as females and different age groups.

Despite these limitations, our study is the first to demonstrate an association between physical activity, prosocial behaviour, and neural markers of intra-brain functional connectivity and inter-brain synchrony. This research should serve as a cornerstone for further investigations into the impact of physical activity on human prosocial behaviour and its neural basis. Based on the present results, future research should consider using face-to-face naturalistic situations and incorporating objectively measured prosocial behaviour, such as economic games, which theoretically share neural mechanisms with the impact of physical activity. Furthermore, exploring the modality (e.g. type, intensity, and duration) of physical activity which can be strongly associated with human prosocial behaviour is a future challenge.

In summary, our findings suggest that physical activity may not merely exhibit a positive correlation with prosocial behaviour; rather, it contributes to the development of adaptable prosocial decision-making in response to social factors. Intra-brain functional connectivity in the DLPFC and inter-brain synchrony in the VLPFC may serve as neural markers of this relationship. Physical activity appears to be associated with greater reciprocity through cognitive control and the sharing of emotions when interacting with others. Furthermore, it seems to be connected to reduced impulsive trust owing to the deliberate consideration of the potential risks of exploitation. Based on the present findings, global trends towards a sedentary lifestyle may be adversely associated with adaptable prosocial decision-making in response to social factors. Consequently, it is imperative to consider the detrimental consequences of the prevalent sedentary lifestyle, not solely from the traditionally emphasized perspective of health risks but also from the perspective of human prosocial behaviour.

## Methods

### Participants

Forty male university students aged 19–24 years were recruited. Demographics of the participants are summarised in Table S4. It is recognised that there are age and sex differences in the impact of various forms of physical activity on cognitive control,^17^ prosocial behaviour during the trust game,^67,68^ and intra-brain functional connectivity.^69^ Considering that this study represents an initial exploration of the correlation between physical activity, prosocial behaviour, intra-brain functional connectivity, and inter-brain synchrony, male participants were chosen exclusively to minimise any potential confounding associations. Eligible participants were required to be free from a history of orthopaedic, cardiovascular, endocrine–metabolic, or visual impairment, mental illness, or other diseases. Furthermore, half of the eligible participants had exercise habits adopted in the Basic Direction for Comprehensive Implementation of National Health Promotion (Health Japan 21, second edition) (≥ 2 times/week, ≥ 30 min/session, over a year or more),^70^ and the rest were not required to have exercise habits (for the reason, see the next paragraph). Before the experiment, all participants completed a pre-screening online questionnaire on demographics (birthday, academic year, and department affiliation), exercise habits, and physical activity. The participants were instructed to avoid consuming alcohol or alcohol-containing products from the day before the experiment. Additionally, on the day of the experiment, the participants were instructed to refrain from consuming caffeine and engaging in exercise. They were instructed to sleep the night before the experiment. This study was approved by the Human Ethics Committee of the Graduate School of Human Development and Environment at Kobe University and all participants provided written informed consent.

To investigate the correlation between physical activity and inter-brain synchronisation, we matched participants into 20 pairs based on their exercise habits (i.e., participants with exercise habits were paired together, and participants without exercise habits were paired together). The relationship between physical activity and inter-brain synchronisation could not be tested by pairing of participants with greater and poor physical activity, due to similar mean inter-brain connectivity within the pair. The pairing criteria were not disclosed to the participants as it could have affected the results.

### Exercise habits and physical activity

Data on exercise habits and physical activity were collected using questionnaires.^54^ Participants provided information on the duration of engagement in exercise, the number of sessions per week, and the duration of each individual session. The frequency and duration were reported separately for weekdays and weekends. Data concerning nine variables were collected, including exercise habits (binary, 0 indicating no exercise and 1 indicating exercise), exercise duration in months, exercise frequency, session duration in minutes, and total weekly exercise duration in minutes. Frequency, session duration, and total weekly duration were recorded separately for weekdays and weekends. Nine data points were included in this analysis (Table S2).

The Japanese version of the International Physical Activity Questionnaire (IPAQ), which has good reliability and validity for assessing physical activity,^55,56^ was used to collect information on various types of physical activity and sedentary behaviours over the preceding week. The IPAQ consists of 31 items and covers activities such as household and yard work, occupational activities, self-powered transport, leisure-time physical activities, and sedentary activities. We calculated the frequency, duration per day, and amount of physical activity (metabolic equivalents × min/week) for each type of activity. Data were analysed separately by category, including walking and moderate- and vigorous-intensity physical activity, and sedentary activities. A total of 46 data points were included in the analysis (Table S2).

### Laboratory procedure

Two experimental conditions were investigated: face-to-face and facial stimulus (Fig. 1a). After visiting the laboratory, the height and weight of the participants were measured. The fNIRS equipment was then worn by them and they were engaged in a practice trial of the trust game until they fully understood the game. To evaluate resting-state functional connectivity, the participants were seated comfortably and instructed to fix their gaze on a fixation cross displayed on a white screen at a distance of approximately 75 cm for 10 min. The visual angle for the fixation cross was subtended by 1.52° horizontally and 1.52° vertically. Following the measurement of resting-state functional connectivity, participants completed the face-to-face and face stimulus conditions in a randomised and counterbalanced order across dyads. During these conditions, the participants engaged in 10 min of eye-to-eye contact with their partner (face-to-face condition) or with a face stimulus displayed on the screen (face stimulus condition). The face stimulus was selected from a face database,^71^ and the chosen face belonged to a male displaying a neutral facial expression. Both conditions were held at consistent distances of approximately 150 and 75 cm, providing a similar visual angle for the face at 6.09° horizontally and 9.09° vertically. Subsequently, the participants completed a trust game with their partners (face-to-face condition) or with a person displayed on the screen (face stimulus condition). A 10-minute break was set before the commencement of both conditions and trust games to allow fNIRS signals to return to baseline levels. After evaluating resting-state functional connectivity and eye-to-eye contact, subjective sleepiness was assessed using a 7-point scale. The participants were instructed to refrain from speaking at any point during the experiment. A partition measuring 190 cm in height and 5 cm in thickness was placed between the participants, except during eye-to-eye contact with their partner, to prevent visual interactions between them. A questionnaire was used at end of the experiment to ensure that the members of each dyad did not know each other.

### Trust game

The trust game is an experiment in which two people, a trustor and trustee, participate. The trustor is given 1000 JPY; he must decide the amount of money that he wishes to give to the trustee in increments of 100 JPY. The amount given is then tripled, and the trustee must decide how to divide the money between him/herself and the trustor. The trustee’s decisions are analysed using the strategy method, which examines the trustee’s response to different amounts given by the trustor in 10% increments. Both participants first make decisions as trustors, and then as trustees. The game ended when a decision was made as a trustee. During the trust game in the face-to-face condition, a partition separated the participants such that they could not see each other’s decisions. The collected data consisted of information on the amount of money transferred for the initial choice and final decisions, as well as the response time for both. The trust game allowed the participants to change their decisions numerous times until the final decision button was pressed. Since intuitive and deliberate decision-making are thought to involve different cognitive processes,^30,61^ both initial and final decisions were included in the analysis. In total, 37 data items related to prosocial behaviour were included in the analyses (Table S2). One participant’s response-time data were not recorded because of a technical malfunction. The average duration of the trust game was approximately 4 min.

### fNIRS data acquisition

The haemodynamic signals of the dyads were simultaneously acquired using a Spectratech OEG-16 and OEG-16H (Spectratech Inc., Tokyo, Japan) CW-NIRS device with 16 active channels (six laser diode emitters, λ1|2 = 840|770 nm with an average power < 5 mW, and six avalanche photodiode detectors) sampled at 0.76 Hz. The probe sets were symmetrically positioned on the scalp over the participant’s prefrontal region, with the centre of each probe located at Fpz, in accordance with the International 10–20 System (Fig. 1c). Each channel consisted of one emitter and one detector placed 3 cm apart (Fig. 1c). The participants were instructed to minimise any head and face movements during the fNIRS measurements. The data for a single channel (channel 1) of one participant were removed owing to a technical malfunction.

### fNIRS data processing and analyses

The fNIRS data were analysed using the OEG-16 software V3.0 (Spectratech Inc., Tokyo, Japan) and R Studio (version 2022.12.0+353). Data were converted to concentration changes in oxy-Hb and deoxy-Hb using the modified Beer–Lambert law^72^ after applying a band-pass filter with cut-off frequencies of 0.001–0.2 Hz (Fig. 1b). After conversion, the fNIRS signals were separated into functional (related to brain activation) and systematic (related to physiological noise) components using the method proposed by Yamada et al. (2012) (Fig. 1b). This method utilises the known characteristics of fNIRS signals in which oxy-Hb and deoxy-Hb are negatively correlated with cerebral function, whereas they are positively correlated with systemic function.^73^ The functional components were used for subsequent analyses.

To evaluate the intra-brain functional connectivity and inter-brain synchrony, two sets of time-series wavelet coherences were computed using *wtc* function in *biwavelet* package. Wavelet coherence is a well-known method for measuring the cross-correlation between two time series as functions of frequency and time.^74^ To calculate the intra-brain functional connectivity, the 16 channels of each participant’s brain were combined with all other channels of the same participant. This resulted in the creation of 120 wavelet coherence maps (16 × 15 channels/2). Intra-brain connectivity data were analysed separately based on intra- and inter-hemispheric connectivity. To calculate the inter-brain synchrony, the 16 channels of each participant’s brain were combined with the 16 channels of their partner’s brain. This resulted in the creation of 256 (16 × 16) wavelet coherence maps. Representative wavelet coherence maps for intra-brain functional connectivity and inter-brain synchrony are shown in Figure 1d. The mean coherence was calculated for two frequency bands, low-frequency band 1 (LF1: 0.015–0.15 Hz) and low-frequency band 2 (LF2: 0.08–0.15 Hz), which have been previously demonstrated to be associated with inter- and intra-brain functional connectivity.^39^ In this procedure, the coherences obtained during the 10-minute breaks that occurred before the start of both conditions and the trust game were excluded. The coherence data were organised into matrices for each experimental condition (resting, control, and face-to-face), activity (eye contact and trust game), and frequency band (LF1 and LF2) (Fig. 1e).

### Confounding variables

To assess the participants’ social value orientation, which reflects stable personality traits towards prosocial behaviour, the slider measure was used,^75^ which is a questionnaire consisting of 15 items that measure the participants’ preference to divide resources between themselves and another person and distinguish between common social value orientations, such as altruistic, prosocial, individualistic, and competitive.^75^ This measure has a good test–retest reliability of r = 0.915.^75^ To assess sleepiness during rest and eye-to-eye contact, a 7-point scale questionnaire was used, which had the following options:1. Energetic, active, clear-headed, fully awake; 2. Awake but not in the best state, able to concentrate on tasks; 3. Relaxed, moderately awake, and able to respond to things; 4. Slightly foggy-headed, feeling like lying down, 5. Dazed, easily distracted, and difficulty staying awake; 6. Sleepy, wanting to lie down, feeling hazy; and 7. Drowsy, unable to stay awake, and likely to fall asleep quickly.

### Statistical analyses

All statistical analyses were conducted using R Studio (version 2022.12.0+353). Prior to the statistical analyses, missing data for the trust game decision-making time (*n* = 1) and fNIRS data (*n* = 1, from a single channel) were filled in using the non-parametric missing value method of the random forest plot through the *missForest* function in the *missForest* package. To test the differences in prosocial behaviour, intra-brain functional connectivity, and inter-brain synchrony between conditions, ANOVAs were performed using the *anova_test* function. Where appropriate, corrections of degrees of freedom for within-participant factors were performed using the Greenhouse–Geisser or Hyunh–Feldt corrections with the *get_anova_table* function. SMCCA was performed using the *MultiCCA.permute* and *MultiCCA* functions in the *PMA* package to test the association between physical activity, prosocial behaviour, intra-brain functional connectivity, and inter-brain synchrony. The SMCCAs were performed separately according to the condition (i.e., face-to-face and face stimulus conditions). Optimal weights and penalties were identified using the *MultiCCA.permute* function with 1,000 permutations. A dimensionality reduction step was performed to prevent an overdetermined, rank-deficient solution and to eliminate the possibility of overfitting. Before performing SMCCA, PCA was performed to reduce the dimensions of all sets of variables using the *prcomp* function. Physical activity and prosocial behaviour data were reduced to 10 PCAs (variance explained = 88% and 95% for both face-to-face and face stimulus conditions, respectively), while intra-brain functional connectivity and inter-brain synchrony data were reduced to 20 PCAs (variance explained = 86% and 73% for the face-to-face condition and 76% and 73% for the face stimulus condition, respectively).

## Data availability

Data supporting the findings of this study are available from the corresponding author upon request.

## Code availability

The scripts for data analysis written in R are available from the corresponding author upon reasonable request.

## Acknowledgments

We thank Yoshiharu Inou, Qiulu Shou, and Kohei Masumoto for their help in conducting this study.

## Authorship contributions

Toru Ishihara: Conceptualization, Data curation, Funding acquisition, Investigation, Methodology, Project administration, Resources, Formal analysis, Visualization, Writing - original draft.

Shinnosuke Hashimoto: Data curation, Investigation. Natsuki Tamba: Data curation, Investigation.

Kazuki Hyodo: Methodology, Writing - review & editing. Tetsuya Matsuda: Writing - review & editing.

Haruto Takagishi: Conceptualization, Methodology, Writing - review & editing.

