## Supplementary information for "Uncovering the links between physical activity and prosocial behaviour: A functional near-infrared spectroscopy hyperscanning study on brain connectivity and synchrony"

**Table S1. The results of ANOVA for prosocial behaviour during trust game**

| Dependent variable | Independent variables | DFn | DFd | F | *p* | Partial $\eta^2$ |
|---|---|---|---|---|---|---|
| Trust | Condition (face stimulus = 0; face-to-face = 1) | 1.00 | 39.00 | 2.63 | 0.11 | 0.06 |
| | Time (final = 0, initial = 1) | 1.00 | 39.00 | 0.39 | 0.53 | 0.01 |
| | Condition × time | 1.00 | 39.00 | 0.66 | 0.42 | 0.02 |
| Reciprocity | Condition (face stimulus = 0; face-to-face = 1) | 1.00 | 39.00 | 9.28 | 0.004 | 0.19 |
| | Time (final = 0, initial = 1) | 1.00 | 39.00 | 3.16 | 0.08 | 0.08 |
| | Amount received (JPY 100 to 1000) | 1.29 | 50.41 | 47.54 | < 0.001 | 0.55 |
| | Condition × time | 1.00 | 39.00 | 0.13 | 0.72 | 0.003 |
| | Condition × amount received | 2.89 | 112.72 | 1.13 | 0.34 | 0.03 |
| | Amount received × time | 3.50 | 136.46 | 2.02 | 0.10 | 0.05 |
| | Condition × time × amount received | 5.28 | 205.96 | 0.60 | 0.71 | 0.02 |
| Decision time | | | | | | |
| Trust (final) | Condition (face stimulus = 0; face-to-face = 1) | 1.00 | 39.00 | 1.57 | 0.22 | 0.04 |
| Trust (initial) | Condition (face stimulus = 0; face-to-face = 1) | 1.00 | 39.00 | 2.59 | 0.12 | 0.06 |
| Reciprocity (final) | Condition (face stimulus = 0; face-to-face = 1) | 1.00 | 39.00 | 9.92 | 0.003 | 0.20 |
| Reciprocity (initial) | Condition (face stimulus = 0; face-to-face = 1) | 1.00 | 39.00 | 2.82 | 0.10 | 0.07 |
| | Amount received (JPY 100 to 1000) | 3.65 | 142.51 | 8.29 | < 0.001 | 0.18 |
| | Condition × amount received | 4.86 | 189.42 | 0.85 | 0.52 | 0.02 |

**Table S2. The list of the measures included in the analyses**

| Physical activity (55 measures) | | |
|---|---|---|
| Regular exercise | Leisure time | Job |
| 1. Exercise habits | 10. Frequency (walk) | 20. Frequency (walk) |
| 2. Duration (year) | 11. Min/day (walk) | 21. Min/day (walk) |
| 3. Frequency (weekday) | 12. MET-min/week (walk) | 22. MET-min/week (walk) |
| 4. Min/day (weekday) | 13. Frequency (MPA) | 23. Frequency (MPA) |
| 5. Frequency (weekend) | 14. Min/day (MPA) | 24. Min/day (MPA) |
| 6. Min/day (weekend) | 15. MET-min/week (MPA) | 25. MET-min/week (MPA) |
| 7. Amount per week (weekday) | 16. Frequency (VPA) | 26. Frequency (VPA) |
| 8. Amount per week (weekend) | 17. Min/day (VPA) | 27. Min/day (VPA) |
| 9. Amount per week (weekend) | 18. MET-min/week (VPA) | 28. MET-min/week (VPA) |
| | 19. MET-min/week (total) | 29. MET-min/week (total) |
| Domestic garden | Sedentary time | Total physical activity |
| 30. Frequency (walk) | 49. Min/day (weekday) | 52. MET-min/week (walk) |
| 31. Min/day (walk) | 50. Min/day (weekend) | 53. MET-min/week (MPA) |
| 32. MET-min/week (walk) | 51. Min/day (total) | 54. MET-min/week (VPA) |
| 33. Frequency (MPA) | | 55. MET-min/week (total) |
| 34. Min/day (MPA) | | |
| 35. MET-min/week (MPA) | | |
| 36. Frequency (VPA) | | |
| 37. Min/day (VPA) | | |
| 38. MET-min/week (VPA) | | |
| 39. MET-min/week (total) | | |

| Prosocial behaviour (35 measures) | | |
|---|---|---|
| Reciprocity (initial choice) | Reciprocity (final decision) | Decision time (reciprocity: initial) |
| 1. 100 JPY | 11. 100 JPY | 21. 100 JPY |
| 2. 200 JPY | 12. 200 JPY | 22. 200 JPY |
| 3. 300 JPY | 13. 300 JPY | 23. 300 JPY |
| 4. 400 JPY | 14. 400 JPY | 24. 400 JPY |
| 5. 500 JPY | 15. 500 JPY | 25. 500 JPY |
| 6. 600 JPY | 16. 600 JPY | 26. 600 JPY |
| 7. 700 JPY | 17. 700 JPY | 27. 700 JPY |
| 8. 800 JPY | 18. 800 JPY | 28. 800 JPY |
| 9. 900 JPY | 19. 900 JPY | 29. 900 JPY |
| 10. 1000 JPY | 20. 1000 JPY | 30. 1000 JPY |
| Decision time (reciprocity: final) | Trust | Decision time (trust) |
| 31. Final decision | 32. Initial choice | 34. Initial choice |
| | 33. Final decision | 35. Final decision |

**Table S3. Participant demographics**

| Variables | Mean | Standard deviation |
|---|---|---|
| Age (years) | 22 | (0.2) |
| Height (cm) | 173.6 | (0.9) |
| Weight (kg) | 69.5 | (1.7) |
| BMI | 23.0 | (0.5) |
| Regular exercise | | |
| Months of exercise (months) | 52.2 | (10.6) |
| Weekday | | |
| Frequency (session/week) | 1.5 | (0.3) |
| Duration (min/session) | 73.5 | (13.2) |
| Total (min/week) | 226.5 | (48.5) |
| Weekend | | |
| Frequency (session/week) | 0.9 | (0.2) |
| Duration (min/session) | 104.5 | (20.2) |
| Total (min/week) | 196.3 | (40.2) |
| Physical activity (MET-minutes/week) | | |
| Overall | | |
| Vigorous intensity | 1542.0 | (350.1) |
| Moderate intensity | 1487.0 | (282.1) |
| Walk | 952.9 | (144.5) |
| Total | 3981.9 | (562.9) |
| At work or university | | |
| Vigorous intensity | 1038.0 | (317.6) |
| Moderate intensity | 499.0 | (113.6) |
| Walk | 322.6 | (79.1) |
| Total | 1859.6 | (379.1) |
| Transportation | | |
| Cycle | 261.0 | (89.5) |
| Walk | 561.0 | (115.2) |
| Total | 822.0 | (139.0) |
| Domestic and garden | | |
| Vigorous intensity (yard chores) | 33.0 | (33.0) |
| Moderate intensity (yard chores) | 38.0 | (30.4) |
| Moderate intensity (inside chores) | 75.0 | (37.6) |
| Total | 146.0 | (73.9) |
| Leisure | | |
| Vigorous intensity | 504.0 | (150.6) |
| Moderate intensity | 581.0 | (174.8) |
| Walk | 69.3 | (28.2) |
| Total | 1154.3 | (252.3) |
| Sedentary time (min/day) | | |
| Weekday | 351.8 | (36.6) |
| Weekend | 342.8 | (39.1) |
| Total | 694.5 | (72.5) |

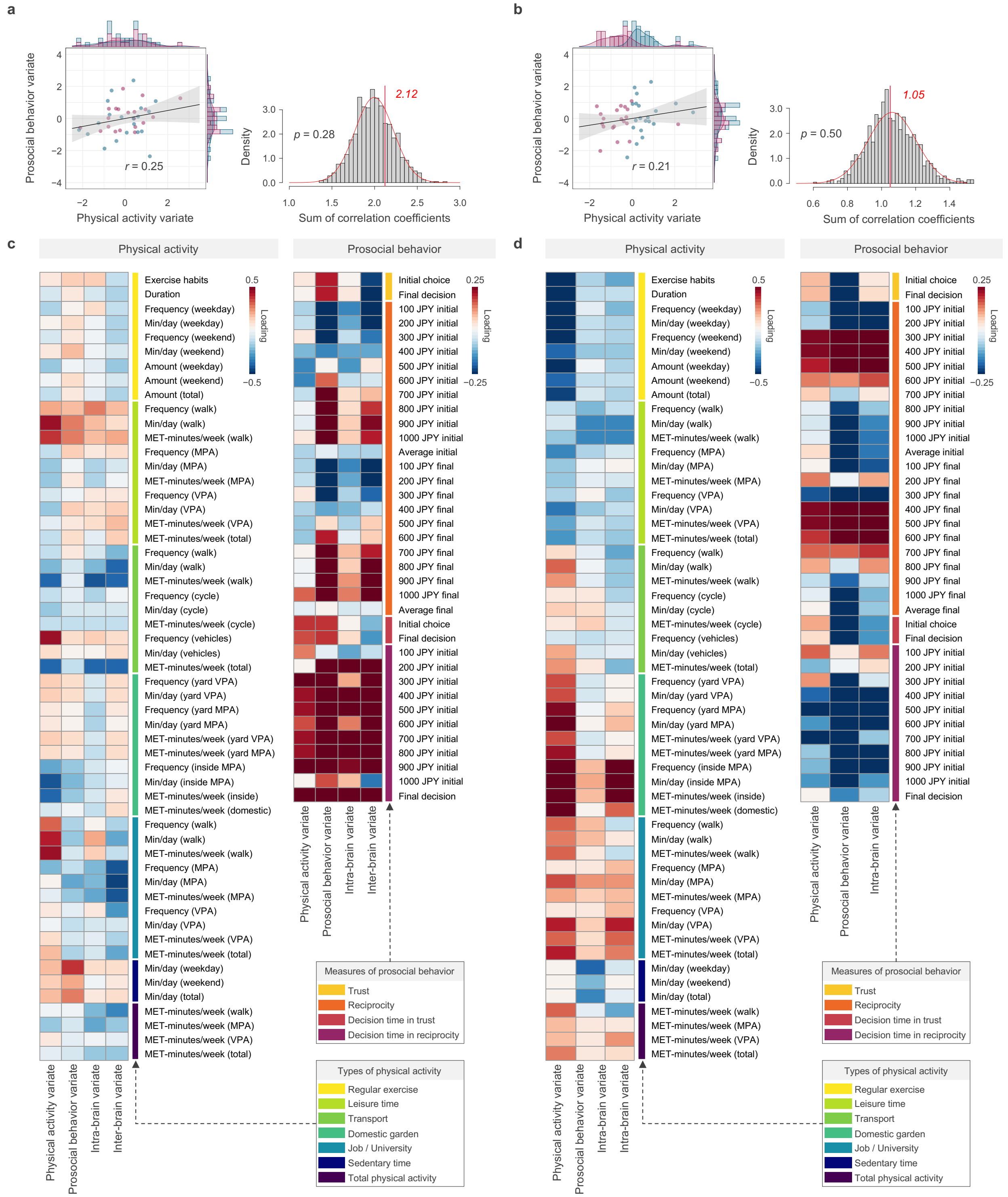


**Fig. S1 | Results of SMCCA during the face stimulus condition.** a,b. The principal SMCCA mode, represented as a scatter plot of physical activity and prosocial behaviours with one point per participant are presented separately by the original model (a) and inter-brain synchrony excluded model (b). An example physical activity measure (exercise habits) is indicated by different colors. c,d. The correlation of physical activity and prosocial behaviour measures with the identified variates are presented separately by the original model (c) and inter-brain synchrony excluded model (d).

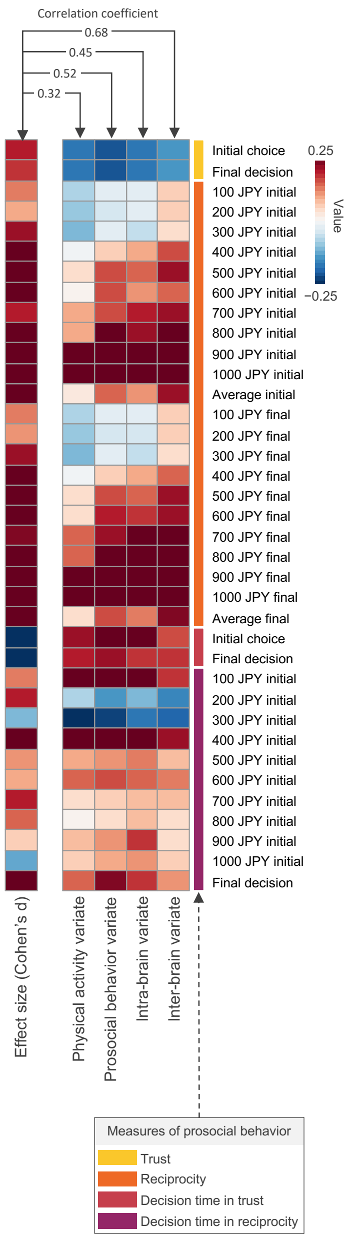


**Fig. S2 | Correlations of differences in prosocial behaviour between conditions (Cohen's d) and canonical loadings.**

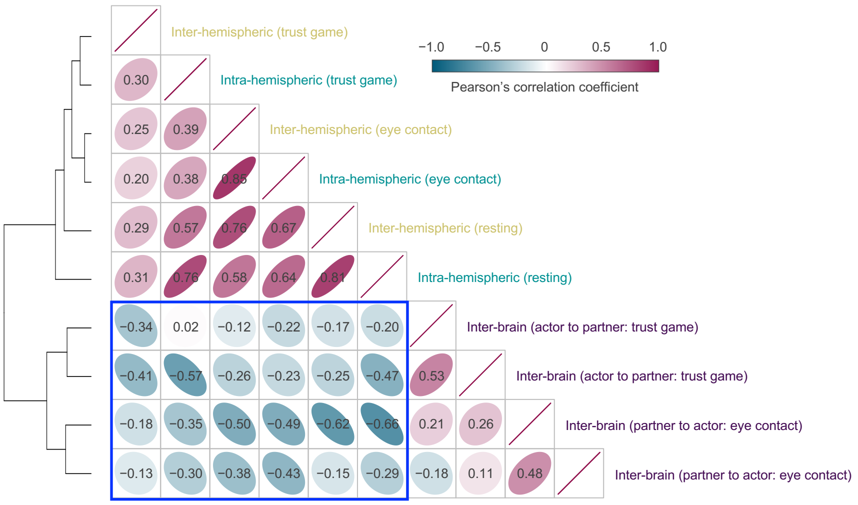


**Fig. S3 | Regional correlations among the SMCCA mode intra-brain functional connectivity and inter-brain synchrony average cross-loadings for physical activity and prosocial behaviour variates.**

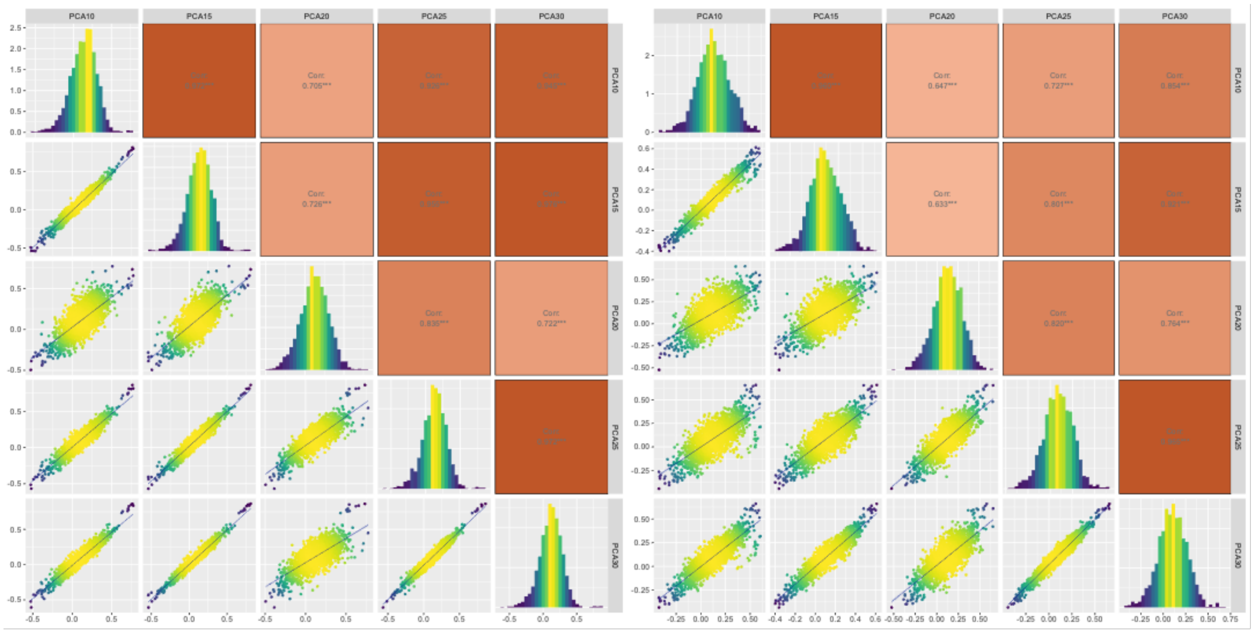


**Fig. S4 | Correlation among canonical- and cross-loadings for physical activity (left panel) and prosocial behaviour (right panel) variates with variation in the number of principal components.**

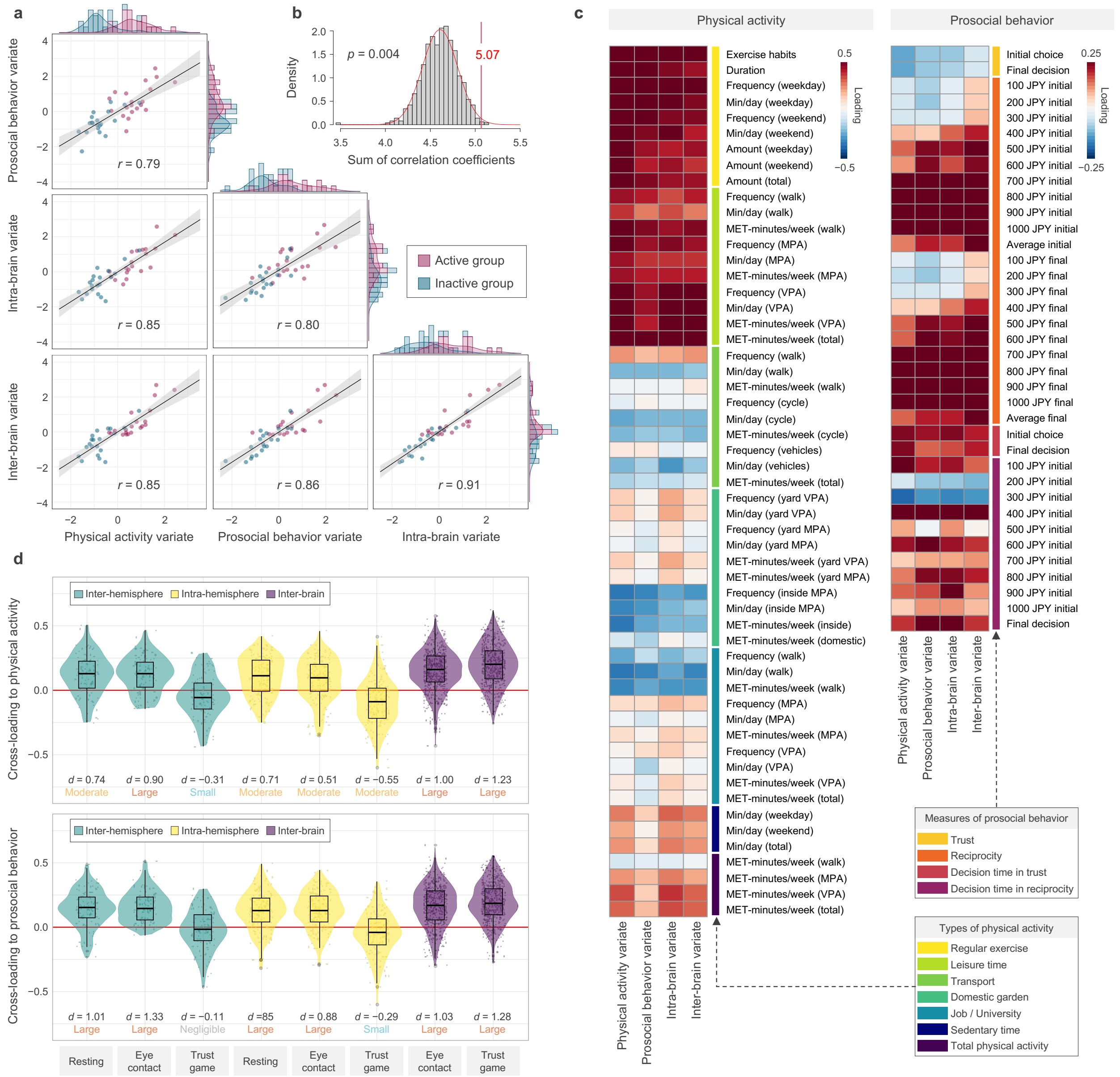


**Fig. S5 | Results of SMCCA during the face-to-face condition adjusted for social value orientation and sleepiness.** a. Principal SMCCA mode, represented as a scatter plot of physical activity, prosocial behaviour, intra-brain functional connectivity, and inter-brain synchrony variates, with one point per participant. An example of physical activity measure (exercise habits) is indicated by different colors. b, Results of the non-parametric permutation test with 1000 iterations. c, The correlation of physical activity and prosocial behaviour measures with the identified variates. d. The correlation of intra-brain functional connectivity and inter-brain synchrony measures with the physical activity (upper panel) and prosocial behaviour (lower panel) variates.

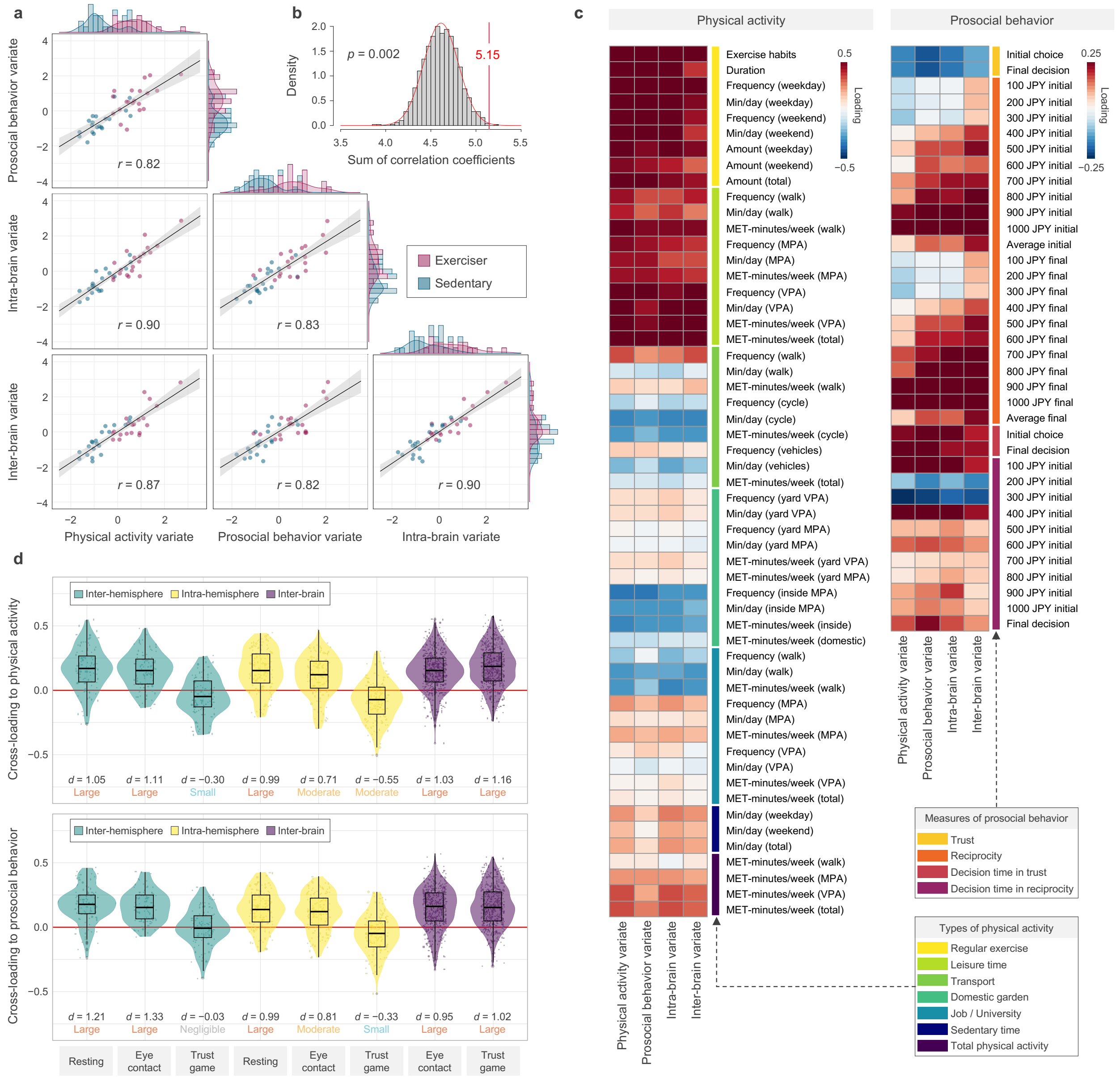


**Fig. S6 | Results of SMCCA during the face-to-face condition using deoxy Hb data.** a. Principal SMCCA mode, represented as a scatter plot of physical activity, prosocial behaviour, intra-brain functional connectivity, and inter-brain synchrony variates, with one point per participant. An example of physical activity measure (exercise habits) is indicated by different colors. b, Results of the non-parametric permutation test with 1000 iterations. c, The correlation of physical activity and prosocial behaviour measures with the identified variates. d. The correlation of intra-brain functional connectivity and inter-brain synchrony measures with the physical activity (upper panel) and prosocial behaviour (lower panel) variates.
